# Heterogeneity Within the Frontoparietal Control Network and its Relationship to the Default and Dorsal Attention Networks

**DOI:** 10.1101/178863

**Authors:** Matthew L Dixon, Alejandro De La Vega, Caitlin Mills, Jessica Andrews-Hanna, R. Nathan Spreng, Michael W. Cole, Kalina Christoff

## Abstract

The frontoparietal control network (FPCN) plays a central role in executive control. It has been predominantly viewed as a unitary domain general system. Here, we examined patterns of FPCN functional connectivity (FC) across multiple conditions of varying cognitive demands, in order to test for FPCN heterogeneity. We identified two distinct subsystems within the FPCN based on hierarchical clustering and machine learning classification analyses of within-FPCN FC patterns. These two FPCN subsystems exhibited distinct patterns of FC the default network (DN) and the dorsal attentional network (DAN). This 2-fold FPCN differentiation was observed across four independent data sets, across 9 different conditions (rest and 8 tasks), as well as in meta-analytic co-activation patterns. The extent of FPCN differentiation varied across conditions, suggesting flexible adaptation to task demands. These findings reveal a flexible and heterogeneous FPCN organization that may in part emerge from separable DN and DAN processing streams.

## INTRODUCTION

Modern neuroscientific investigations have demonstrated that frontoparietal cortices contribute to executive control and adaptive behavior via the flexible encoding of task demands and desired outcomes, and the top-down modulation of processing in other brain regions^1-8^. Despite this progress, we lack a clear understanding of the functional organization of frontoparietal cortex, a critical step in discerning the network architecture underlying executive control. Distributed frontoparietal regions often activate together in response to diverse task demands, suggesting that they may function as a unified, domain general control system, referred to as the frontoparietal control network (FPCN) or “multiple demand” system^4^. It is possible, however, that a finer-level of organization may be present within the FPCN, with distinct subsystems contributing to different types of executive control. Progress has been made in understanding other networks (e.g., default network) via fractionating them into distinct subsystems with unique functional roles^9^. Existing models have distinguished the FPCN from networks centered on insular and cingulate cortices (e.g., “salience” and cingulo-opercular networks)^10,11^. However, possible heterogeneity within the FPCN has not been explored in detail.

In a seminal paper, Yeo and colleagues^12^ introduced a 7-network parcellation that has had a considerable influence on the field of network neuroscience. In this 7-network parcellation, the FPCN appears as a uniform network. However, Yeo et al. also reported a fine-grained 17-network parcellation that has received much less attention in the literature. In this 17-network solution, the FPCN appears to be segregated into 2 distinct subsystems (see Yeo et al., Figure 9). Recent work suggests that a fractionation of the FPCN can be observed in the data of individual participants^13^. These findings represent important empirical evidence for functional heterogeneity within this network. However, prior work has not systematically investigated the basis for this FPCN fractionation or its functional implications.

Here, we used a hypothesis-driven approach together with graph theoretical analyses to examine the possibility that the fine-grained organziation of the FPCN may be driven by specific connectional patterns as part of a “distance from sensorimotor processing” principle that defines global brain organization^14-17^. The FPCN is extensively interconnected with both the default network (DN) and dorsal attention network (DAN)^18^―large-scale systems that contribute to distinct, and sometimes competing modes of processing^19,20^. The DAN has a close relationship with sensorimotor regions^12^ and plays a key role in visuospatial perceptual attention^21,22^. It contains neurons with spatially organized receptive fields^22,23^ that are activated during saccades^24^, shifts of attention to salient objects in the external environment^25-27^, and during reaching actions towards such objects^21^. In contrast, the DN contributes to social, conceptual, and associative thought processes that are, in some cases, independent from sensory input^14,28-31^. Specifically, the DN is involved in mentalizing^32^, autobiographical memory^33^, spontaneous cognition^34-37^, self-referential processing^38^, and high-level aspects of emotion^29,39^. Correspondingly, it has been demonstrated that the DN is further removed spatially and functionally from sensorimotor processing than is the DAN^14^. Notably, there is also some evidence of a gradient of representational abstraction within parts of the FPCN, specifically, the lateral prefrontal cortex^40-46^. Based on these findings, we hypothesized that the distinct DN and DAN processing streams may be carried forward into the organization and functions of the FPCN. While prior work has documented functional connections linking the FPCN to these networks^18^, here we predicted that these connections may exhibit a specific topographical organization.

We first examined the network architecture of the FPCN using hierarchal clustering to determine whether FPCN nodes separate into distinct subsystems based on intra-modular (within-network) connections. We then determined whether the observed subsystems exhibit topographically organized functional connections with the DN and DAN. That is, we predicted that FPCN regions coupled with the DN would be spatially distinct from FPCN regions coupled with the DAN. We investigated functional coupling patterns during rest and several different tasks, which allowed us to look for differences in coupling patterns that persist across different cognitive states. Second, to determine the generalizability of a putative FPCN fractionation related to the DN and DAN, we examined FC patterns in three additional independent data sets, and examined meta-analytic co-activation patterns across 11,406 neuroimaging studies within the Neurosynth database^47^. Third, we examined the temporal evolution of network properties, and investigated whether dynamic FC patterns also display evidence of a FPCN fractionation. Specifically, we examined whether spatially-specific FPCN interactions correlate with time-varying changes in the capacity for specialized processing within the DN and DAN, indexed with a graph theoretic measure known as the clustering coefficient^48^. We also examined how the FPCN fractionation relates to task-related flexibility in FC patterns. Finally, in an exploratory analysis, we used Neurosynth topic mapping to identify functional domains that differentially predict activation in the subsystems.

Our primary data set involved data collected from 24 participants that underwent fMRI scanning during six separate conditions designed to elicit mental states similar to those frequently experienced in everyday life. These six conditions varied in the amount of abstract conceptual thought and perceptual demands, and included: (i) rest; (ii) movie viewing; (iii) analysis of artwork; (iv) social preference shopping task; (v) evaluation-based introspection; and (vi) acceptance-based introspection (see **Methods** for details). Additionally, we examined FC patterns in three other data sets involving traditional cognitive control tasks that are known to activate the FPCN: (i) rule use; (ii) Stroop; (iii) 2-Back working memory. Data were processed using standard techniques^49^, and we did not use global signal regression, so as to avoid distorting FC values^50^.

## RESULTS

### Evidence for distinct FPCN subsystems

Graph theory represents complex systems such as the brain as a graph consisting of a set of nodes (regions) and edges (connections between nodes), and allows for a quantitative description of network properties^48,51^. We calculated the time-series correlation between nodes spanning the DAN, DN, and FPCN based on the Yeo parcellation^12^. We first analyzed the organization of FPCN nodes based on intra-modular (within-network) FC patterns. We used hierarchical clustering to organize nodes into a tree structure based on the similarity of their FC profiles. The analysis revealed clear evidence of two distinct clusters or subsystems that we refer to as FPCN_A_ and FPCN_B_ (**Fig. 1a-b**). FPCN_A_ and FPCN_B_ regions were somewhat interleaved, similar to observations in prior work^12,13^. To examine whether the distinction between FPCN_A_ and FPCN_B_ FC patterns were consistent across participants, we used a linear support vector machine (SVM) classifier to distinguish FPCN_A_ and FPCN_B_ intra-modular FC patterns in new participants based on data from other participants. The SVM attempts to find a hyperplane that best separates the two classes of data. We used k-fold cross-validation (*k*=4) where the classifier was trained on data from 75% of participants then tested on unlabeled data from the remaining 25% of participants. Using this procedure, we found highly accurate (> 90 %) discrimination of the FPCN_A_ and FPCN_B_ during every condition in the primary data set (**Fig. 1c; Supplementary Fig. 2**). Permutation testing in which FPCN subsystem labels were randomly shuffled revealed chance level discrimination (∼ 50% accuracy; see **Methods**). A FPCN fractionation was also observed when using an independent set of nodes and network definitions based on the Gordon parcellation^52^ (**Fig. 1d**), or Power parcellation^53^ (**Supplementary Fig. 1**).

**Figure 1.**
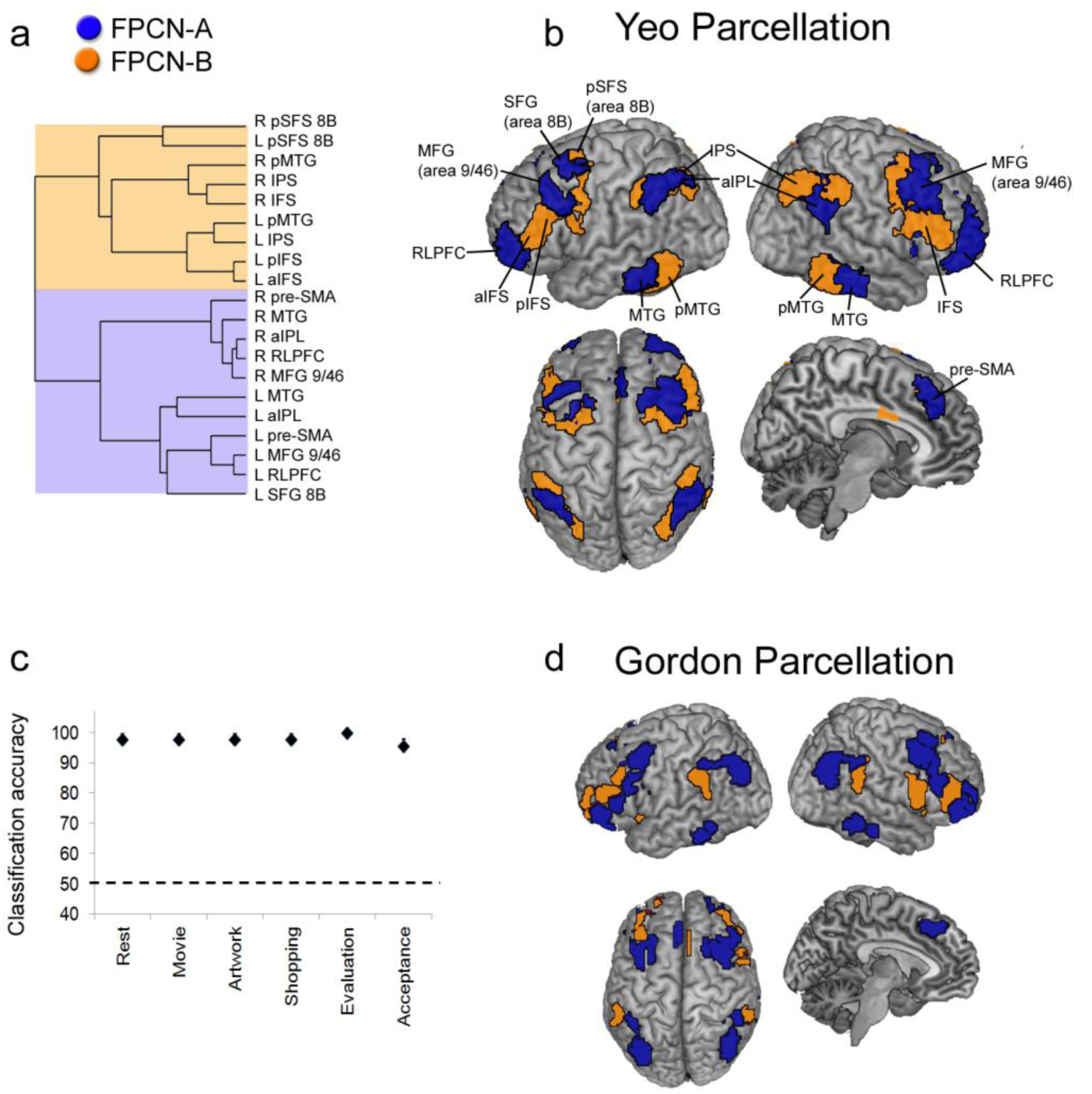
FPCN fractionation based on intra-modular connectivity. **(a)** Hierarchical clustering results based on intra-modular (within-FPCN) connections. FPCN nodes cluster into two separate families. **(b)** Surface rendering of FPCN nodes from the Yeo parcellation, color-coded based on the hierarchical clustering results in (a). **(a)** Accuracy of the support vector machine classifier in distinguishing FPCN_A_ and FPCN_B_ within-network FC patterns during each condition. Dotted line represents baseline accuracy (50%). **(d)** Surface rendering of FPCN nodes from the Gordon parcellation, color-coded based on the hierarchical clustering results in **Supplementary Figure 1a**. Abbreviations: RLPFC, rostrolateral prefrontal cortex; MFG, middle frontal gyrus; aIPL, anterior inferior parietal lobule; MTG, middle temporal gyrus; pre-SMA, pre-supplementary motor area; aIFS, anterior inferior frontal sulcus; pIFS, posterior inferior frontal sulcus; IPS, intraparietal sulcus; pMTG, posterior middle temporal gyrus; pSFS, posterior superior frontal gyrus.

We next examined FPCN clustering patterns based strictly on functional connections with the DN and DAN. The results again revealed two distinct subsystems, nearly identical to the structure observed based on intra-modular connections (**Fig. 2; Supplementary Fig. 3**). The separation between FPCN_A_ and FPCN_B_ based on FC with the DN and DAN was highly consistent across participants, as evidenced by highly accurate discrimination when using a linear SVM classifier and 4-fold cross-validation (**Fig. 2c; Supplementary Fig. 4**). Together, these findings reveal heterogeneity within the FPCN based on intra-modular connections and based on inter-network connectivity patterns with the DN and DAN.

**Figure 2.**
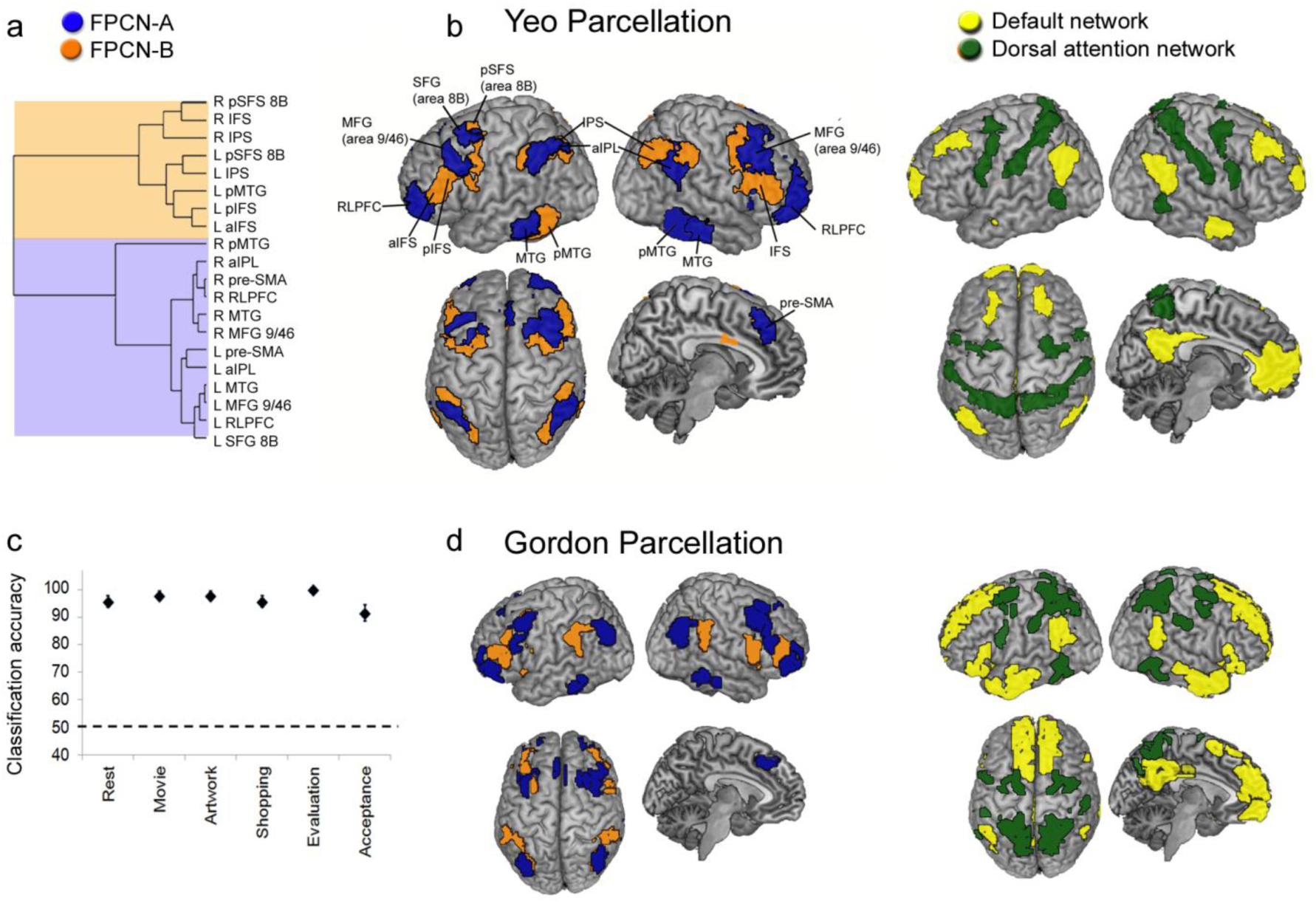
FPCN fractionation based on inter-network connectivity with the DN and DAN. **(a)** Hierarchical clustering results based on inter-modular connections. FPCN nodes cluster into two separate families. **(b)** Surface rendering of FPCN nodes from the Yeo parcellation, color-coded based on the hierarchical clustering results in A. **(c)** Accuracy of the support vector machine classifier in distinguishing FPCN_A_ and FPCN_B_ FC patterns with the DN and DAN during each condition. Dotted line represents baseline accuracy (50%). (d) Surface rendering of FPCN nodes from the Gordon parcellation, color-coded based on the hierarchical clustering results in **Supplementary Figure 3a**. Abbreviations: RLPFC, rostrolateral prefrontal cortex; MFG, middle frontal gyrus; aIPL, anterior inferior parietal lobule; MTG, middle temporal gyrus; pre-SMA, pre-supplementary motor area; aIFS, anterior inferior frontal sulcus; pIFS, posterior inferior frontal sulcus; IPS, intraparietal sulcus; pMTG, posterior middle temporal gyrus; pSFS, posterior superior frontal gyrus.

### Differential coupling patterns with the DN and DAN

To elucidate the underlying basis of the FPCN fractionation, we visualized the network topology using the Kamada–Kawai energy algorithm^54^, which produces spring-embedded layouts that minimize the geometric distances of nodes based on their topological distances in the graph. Nodes are pulled together or pushed apart based on the strength of functional connections rather than anatomical locations. The network visualization revealed a clear separation of FPCN_A_ and FPCN_B_ nodes during all six conditions in the primary data set, with FPCN_A_ preferentially connecting to DN nodes, and FPCN_B_ preferentially connecting to DAN nodes (**Fig. 3**). The group-averaged correlation matrix revealed that FPCN_A_ nodes exhibited positive correlations with DN nodes and no correlation or negative correlations with DAN nodes, whereas FPCN_B_ nodes exhibited the opposite pattern (**Fig. 4a**). Furthermore, FC fingerprints (**Fig. 4b**) and whole-brain seed-based correlation maps (**Supplementary Fig. 5**) revealed that spatially adjacent FPCN_A_ and FPCN_B_ nodes exhibited highly divergent functional coupling patterns with DN and DAN regions. Importantly, differences in FPCN_A_ and FPCN_B_ coupling patterns were not driven by spatial proximity to DN and DAN nodes (**Supplementary Results**). Thus, distinct FPCN subsystems can be delineated based on topographically organized functional connections with the DN and DAN.

**Figure 3.**
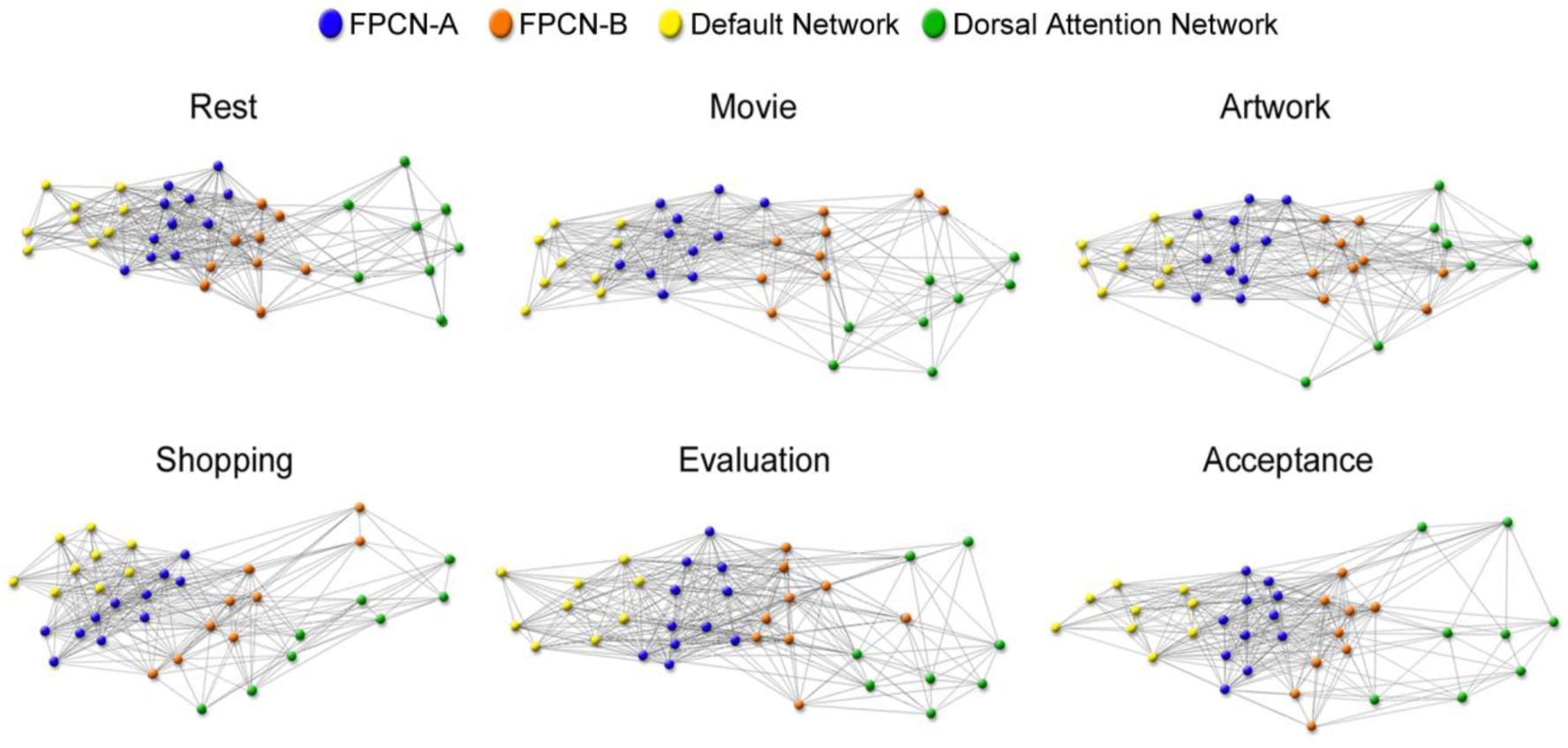
Visualization of the network topology. FPCN nodes are color-coded based on the hierarchical clustering analysis of intra-modular connections using the Yeo parcellation nodes. In every context, there is a clear separation of FPCN nodes, with FPCN_A_ nodes exhibiting preferential FC with DN nodes, and FPCN_B_ nodes exhibiting preferential FC with DAN nodes.

**Figure 4.**
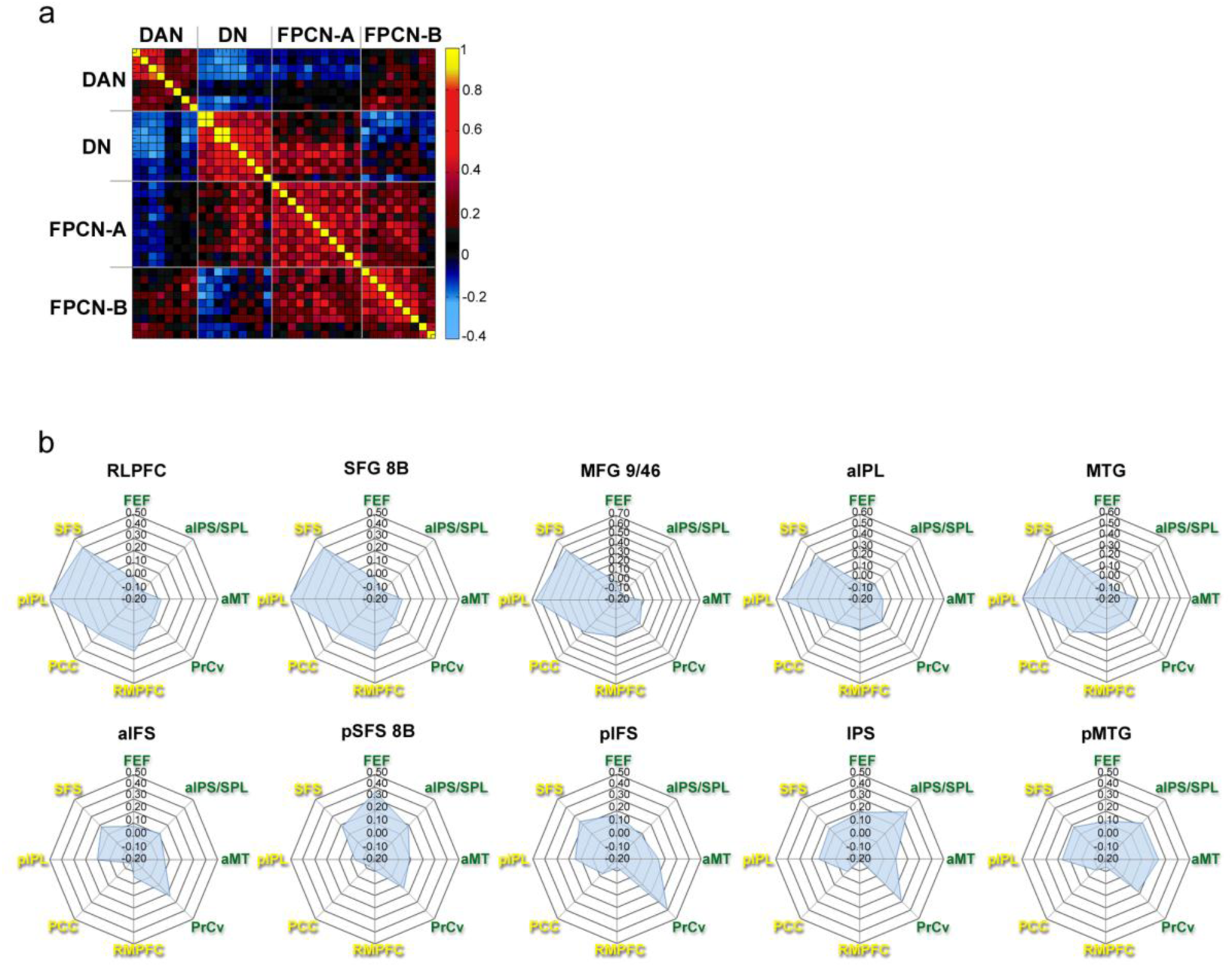
Differential FPCN subsystem coupling patterns. **(a)** Group-averaged correlation matrix reflecting mean *z(r)* values across the six task conditions, using Yeo parcellation nodes. **(b)** FC fingerprints for each FPCN node. Top panel: FPCN_A_ nodes demonstrate a clear leftward bias, reflecting stronger FC with DN nodes (yellow text). Bottom panel: FPCN_B_ show a slight rightward bias reflecting stronger FC with DAN nodes (green text), though there is evidence of FC with DN nodes as well. Critically, FPCN_A_ and FPCN_B_ fingerprints are highly divergent for each pair of spatially adjacent nodes (top versus bottom panel fingerprint). Abbreviations: RLPFC, rostrolateral prefrontal cortex; MFG, middle frontal gyrus; aIPL, anterior inferior parietal lobule; MTG, middle temporal gyrus; pre-SMA, pre-supplementary motor area; aIFS, anterior inferior frontal sulcus; pIFS, posterior inferior frontal sulcus; IPS, intraparietal sulcus; pMTG, posterior middle temporal gyrus; pSFS, posterior superior frontal gyrus; RMPFC, rostromedial prefrontal cortex; PCC, posterior cingulate cortex; pIPL, posterior inferior parietal lobule; SFS, superior frontal suclus; LTC, lateral temporal cortex; FEFs, frontal eye fields; aIPS/SPL, anterior intraparietal sulcus/superior parietal lobule; PrCv, ventral precentral cortex; aMT anterior middle temporal region.

To quantify coupling patterns, we computed the average strength of FC between each pair of networks (**Fig. 5**). A 2 (FPCN_A_ vs FPCN_B_) x 2 (DN vs DAN) repeated measures ANOVA revealed a robust interaction during every condition [all *F*_1, 23_ > 96.83, *P*'s < .001)]. In each case, FPCN_A_-DN coupling was stronger than FPCN_B_-DN coupling (paired *t*-test: all *t*_23_ > 9.01, *P*'s < .05, Bonferroni corrected), whereas FPCN_B_-DAN coupling was stronger than FPCN_A_-DAN coupling (paired *t*-test: all *t*_23_ > 6.93, *P*'s < .05, Bonferroni corrected). While the DN Core subsystem was our main focus, for completeness we also report relationships involving the other subsystems of the DN in **Supplementary Figure 6**. Interestingly, a “selectivity index” (see **Methods**) revealed that differential coupling was stronger for FPCN_A_ nodes than FPCN_B_ nodes [paired *t*-test: *t*_23_ = 4.78, *P* < .001] (**Supplementary Fig. 7**).

**Figure 5.**
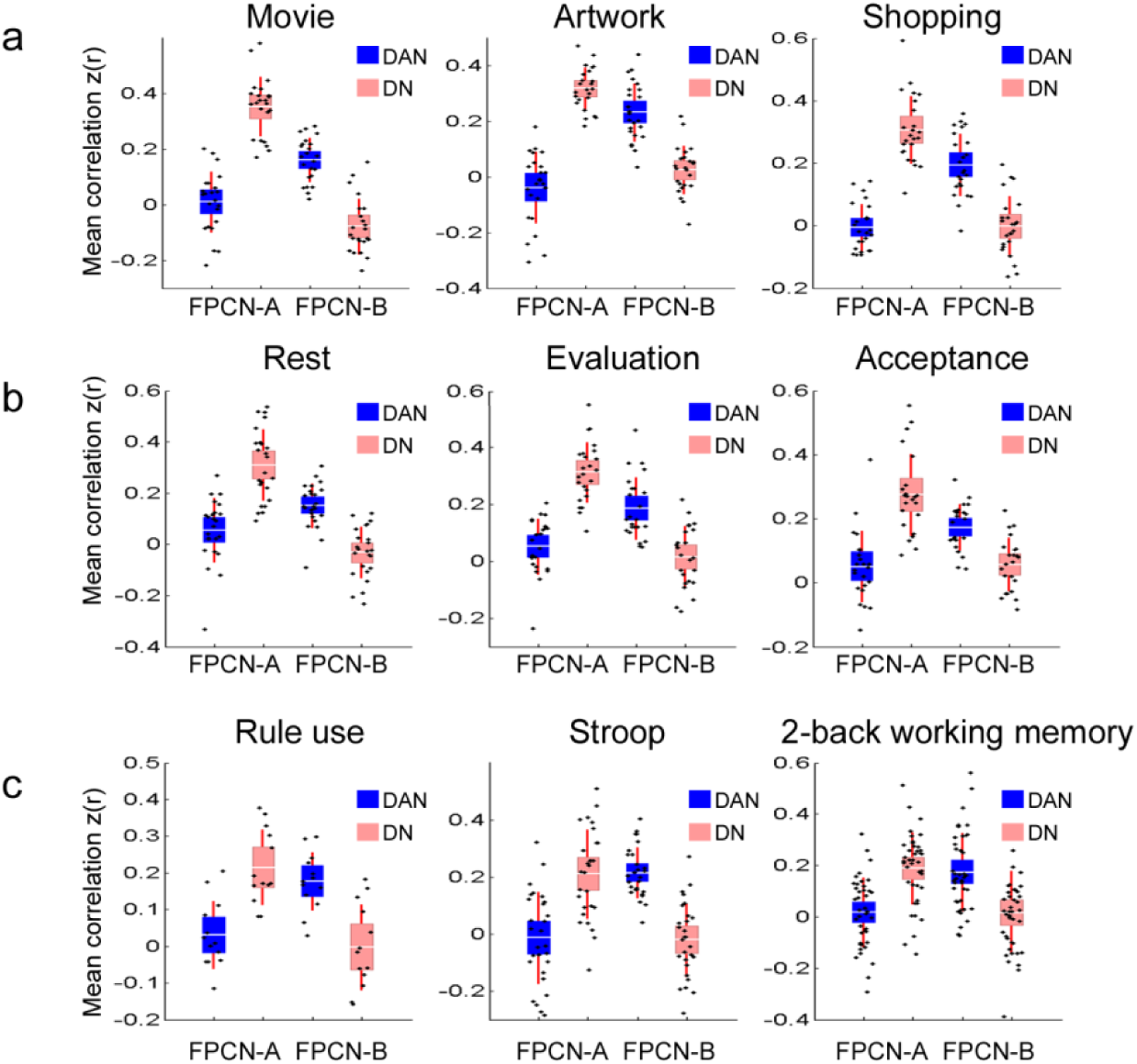
Mean function connectivity between the FPCN subsystems and the DN and DAN, using Yeo parcellation nodes. Conditions are separated into: **(a)** tasks with a perceptual component; **(b)** tasks without a perceptual component; and **(c)** cognitive control tasks from the replication samples. Data for each participant (black dots), with mean (white line), 95% CI (light red and blue shaded areas) and 1 SD (dark red lines).

### Replication and generalizability of differential coupling patterns

We next examined whether the fractionation would replicate in three independent data sets involving demanding cognitive control tasks (rule use; Stroop; 2-Back working memory). We found a robust FPCN subsystem x DN/DAN interaction in each task, with FPCN_A_ preferentially coupling with the DN, and FPCN_B_ preferentially coupling with the DAN [rule use: *F*_1,14_ = 109.84, *P* < .001; Stroop: *F*_*1,27*_ *=* 189.17, *P* < .001; N-Back: *F*_1,40_ = 108.40, *p* < .001] (**Fig. 5c**).

To examine the generalizability of the FPCN fractionation we performed an automated meta-analysis on coactivation patterns across the wide range of tasks within the Neurosynth database^47^. The results demonstrated that there are notable differences in co-activation with other parts of the brain between the two FPCN subsystems, consistent with our predictions (**Fig. 6**). In particular, FPCN_A_ co-activates to a greater extent with the default network (e.g., rostromedial PFC, posterior cingulate cortex, lateral temporal cortex), than does FPCN_B_. There was less evidence for a distinction with respect to co-activation with the DAN. However, FPCN_B_ does co-activate to a greater extent with portions of DAN around the superior parietal lobule and frontal eye fields.

**Figure 6.**
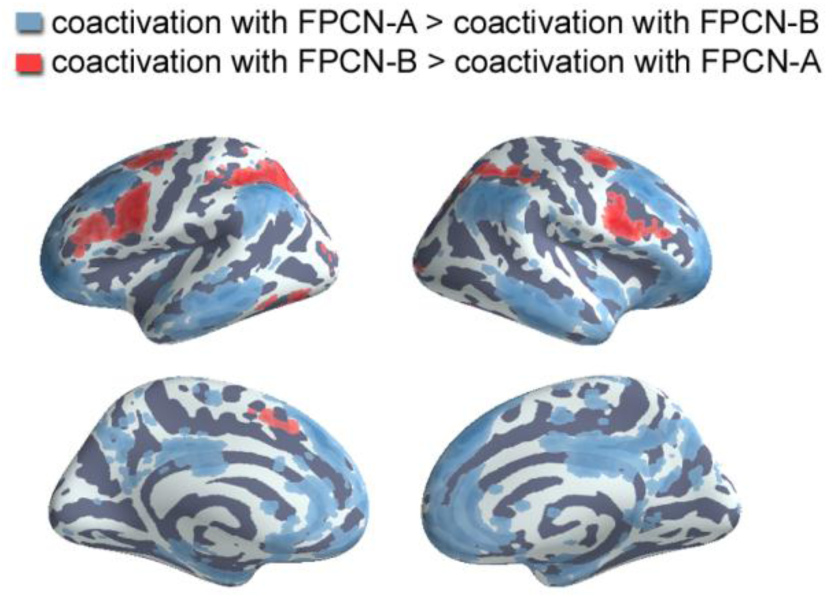
Meta-analytic coactivation contrasts. Red voxels indicate significantly greater coactivation with FPCN_B_ than FPCN_A_. Blue voxels indicate significantly greater coactivation with FPCN_A_ than FPCN_B_. Images were whole-brain corrected using a false discovery rate of *q* = 0.05.

### Dynamic evolution of differential coupling patterns and network clustering

We next asked whether differential coupling is present in dynamic network interactions, and whether these interactions correlate with time-dependent changes in the capacity for specialized processing within the DN and DAN. We quantified the nature of processing within the DN and DAN using a modified version of the clustering coefficient, a graph theoretic measure which computes the proportion of neighbors around node *i* that are also interconnected^48^. When normalized by the connection strength (weight) between nodes, the clustering coefficient provides an index of the strength of communication within a densely-connected neighborhood (which in this case was a specific network; see **Methods**). Using a sliding window approach, we derived a time-series of mean weighted clustering coefficients for the DN and DAN. We also derived a time-series of between-network FC values. We then used the Pearson correlation to quantify the relationship between time-dependent coupling patterns and clustering strength.

As illustrated in **Figure 7**, DN and DAN within-network processing strength (i.e., mean weighted-clustering) varied considerably across time. Critically, these changes were tightly coupled with the strength of interactions involving the FPCN subsystems (**Fig. 7**). Periods of time characterized by stronger FPCN_A_-DN coupling were associated with larger clustering coefficients for the DN, whereas temporal variation in FPCN_B_-DN coupling was unrelated to changes in DN clustering. The relationship between FPCN_A_-DN coupling and DN clustering was significantly stronger than the relationship between FPCN_B_-DN coupling and DN clustering in every condition (paired *t*-test: all *t*'s > 4.17 *P*'s < .05, Bonferroni corrected) with the exception of the acceptance-based introspection condition which did not reach significance (*P* = .26, Bonferroni corrected). On the other hand, periods of time characterized by stronger FPCN_B_-DAN coupling were associated with larger clustering coefficients for the DAN, whereas temporal variation in FPCN_A_-DAN coupling was unrelated to changes in DAN clustering. The relationship between FPCN_B_-DAN coupling and DAN clustering was significantly stronger than the relationship between FPCN_A_-DAN coupling and DAN clustering in every condition (paired *t*-test: all *t*'s > 3.96, *P*'s < .05, Bonferroni corrected) with the exception of the shopping (*P* = .09, Bonferroni corrected) and two introspection conditions (evaluation: *P* = .87, Bonferroni corrected; acceptance: *P* = .14, Bonferroni corrected) which did not reach significance. Control analyses revealed that these patterns were unrelated to motion (**Supplementary Results**). Thus, dynamic interactions between FPCN_A_ and the DN were specifically associated with temporal variation in the strength of communication within the DN, whereas dynamic interactions between FPCN_B_ and the DAN were specifically associated with temporal variation in the strength of communication within the DAN. These findings reveal clear structure in how network properties evolve across time, and reinforce the idea that FPCN organization has a close relationship with the DN and DAN.

**Figure 7.**
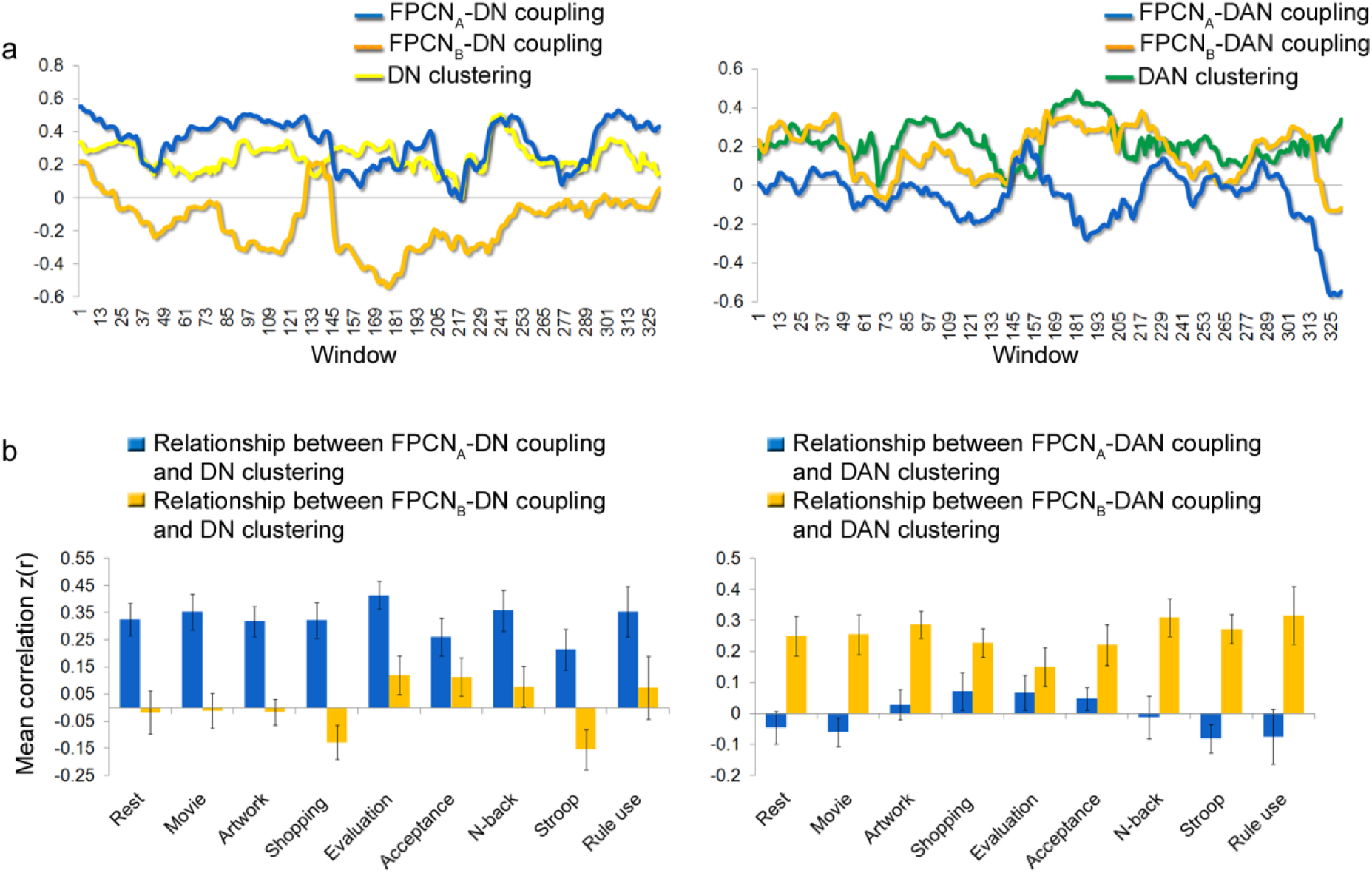
Dynamic network interactions and clustering. **(a)** Example participant data from the Stroop task demonstrating the relationship between temporal fluctuations in FPCN interactions with the DN and DAN and mean weighted clustering strength. **(b)** Mean correlation between changes across time in clustering and between-network FC strength across participants. Fluctuations in FPCN_A_-DN FC are positively correlated with fluctuations in DN clustering strength, and fluctuations in FPCN_B_-DAN FC are positively correlated with fluctuations in DAN clustering strength. Error bars represent between-subject SEM.

### FPCN fractionation and task-related flexibility

Prior work has shown that FPCN FC patterns exhibit a high-level of task-related flexibility^5,7,55^. We examined how differential coupling patterns relate to this type of flexibility. We computed a task-related “flexibility index” reflecting the extent to which FC patterns changed more across conditions than within conditions from the first half to the second half. This measure of flexibility pertains to context and is different from the measure used by Bassett and colleagues which pertains to flexibility in the temporal domain^56^. One sample *t*-tests revealed that both subsystems exhibited a significant flexibility index, revealing task-dependent reconfiguration of FC patterns [FPCN_A_: *t*_23_ = 5.62, *P* < .001; FPCN_B_: *t*_*23*_ = 8.86, *P* < .001] (**Supplementary Fig. 8a**). Interestingly, FPCN_B_ demonstrated stronger task-related flexibility than FPCN_A_ [paired *t*-test: *t*_*23*_ = 2.25, *P* = .043]. Not only did overall FC with the DN and DAN change across conditions for both subsystems, but so did the magnitude of the “selectivity index”―the relative strength of DN to DAN connections (**Supplementary Fig. 8b**). Interestingly, the “selectivity index” was weakest during the traditional cognitive control tasks (**Supplementary Fig. 8b**). Thus, while FPCN_A_ and FPCN_B_ exhibited differential coupling patterns in every condition, the magnitude of this effect was sensitive to task demands. The right IFS/IFJ node of FPCN_B_ exhibited the greatest FC flexibility (**Supplementary Fig. 8c-e**).

### Are FPCN_A_ and FPCN_B_ subsystems of the same network or extensions of the DN and DAN?

To determine whether FPCN_A_ and FPCN_B_ should be considered subsystems within the same network or extensions of the DN and DAN we compared mean between-network and between-subsystem FC patterns using paired *t*-tests. During the traditional cognitive control tasks, FPCN_A_ and FPCN_B_ exhibited stronger coupling with each other than with the DN (rule use: *t*_14_ = 3.21, *P* = .057, Bonferroni corrected; Stroop: *t*_27_ = 6.16, *P* < .001, Bonferroni corrected; 2-Back: *t*_41_ = 4.69, *P* < .001, Bonferroni corrected) or DAN (rule use: *t*_14_ = 7.41, *P* < .001, Bonferroni corrected; Stroop: *t*_27_ = 7.67, *p* < .001, Bonferroni corrected; 2-Back: *t*_41_ = 5.05, *P* < .001, Bonferroni corrected) (**Fig. 8**). However, the picture is less clear during the other conditions that involved a range of processing demands. Coupling between FPCN_A_ and FPCN_B_ was weaker than FPCN_A_-DN coupling during the movie (*t*_22_ = 4.30, *P* < .05, Bonferroni corrected) and shopping conditions (*t*_23_ = 3.27, *P* < .05, Bonferroni corrected) but not different during the other conditions (*P*'s > .05, Bonferroni corrected). Coupling between FPCN_A_ and FPCN_B_ was stronger than FPCN_B_-DAN coupling during rest (*t*_23_ = 9.82, *P* < .05, Bonferroni corrected), evaluation (*t*_23_ = 5.15, *P* < .05, Bonferroni corrected), and acceptance (*t*_23_ = 7.59, *P* < .05, Bonferroni corrected), but not different during the other conditions (*P*'s > .05, Bonferroni corrected). These findings suggest that the extent to which the FPCN_A_ and FPCN_B_ cluster together versus with the DN/DAN depends on current processing demands.

**Figure 8.**
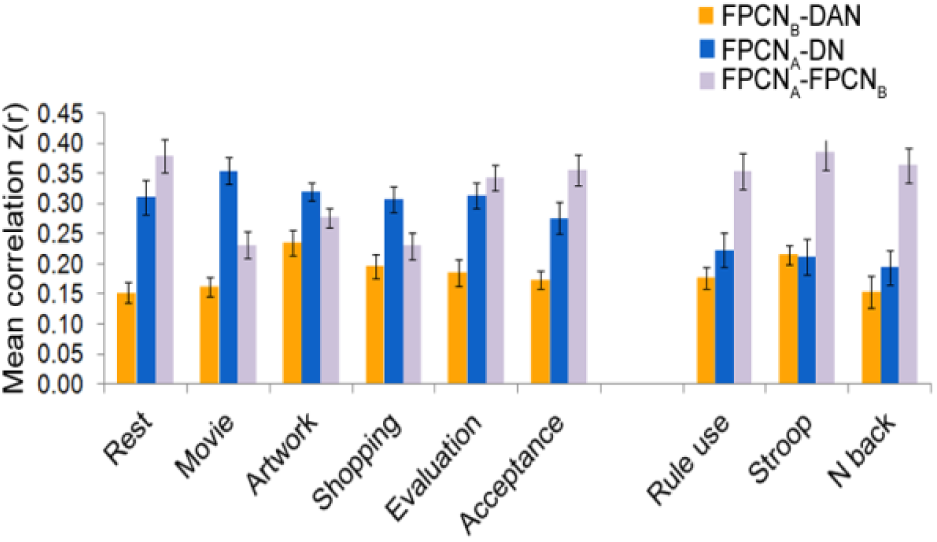
Mean between-network FC in each condition.

### Meta-analytic functional differentiation

To examine whether the FPCN subsystem distinctions in network architecture are functionally meaningful, we used a naive Bayes classifier to determine which Neurosynth topics were preferentially associated with each subsystem. We plotted the loading of each topic onto each subsystem along with bootstrapped 95% confidence intervals (**Fig. 9; Supplementary Fig 9**). As expected, both subsystems showed high loadings to executive function topics including working memory, switching, and conflict. Notably, there were also clear distinctions. The topics “mentalizing” and “emotion” loaded more strongly onto FPCN_A_ than FPCN_B._ In contrast, “attention”, “action”, “reading”, and “semantics” loaded more strongly onto FPCN_B_ than FPCN_A_. These differences are consistent with the idea that FPCN_A_ is biased toward functions that are associated with the DN, whereas FPCN_B_ is biased toward functions that are associated with the DAN.

**Figure 9.**
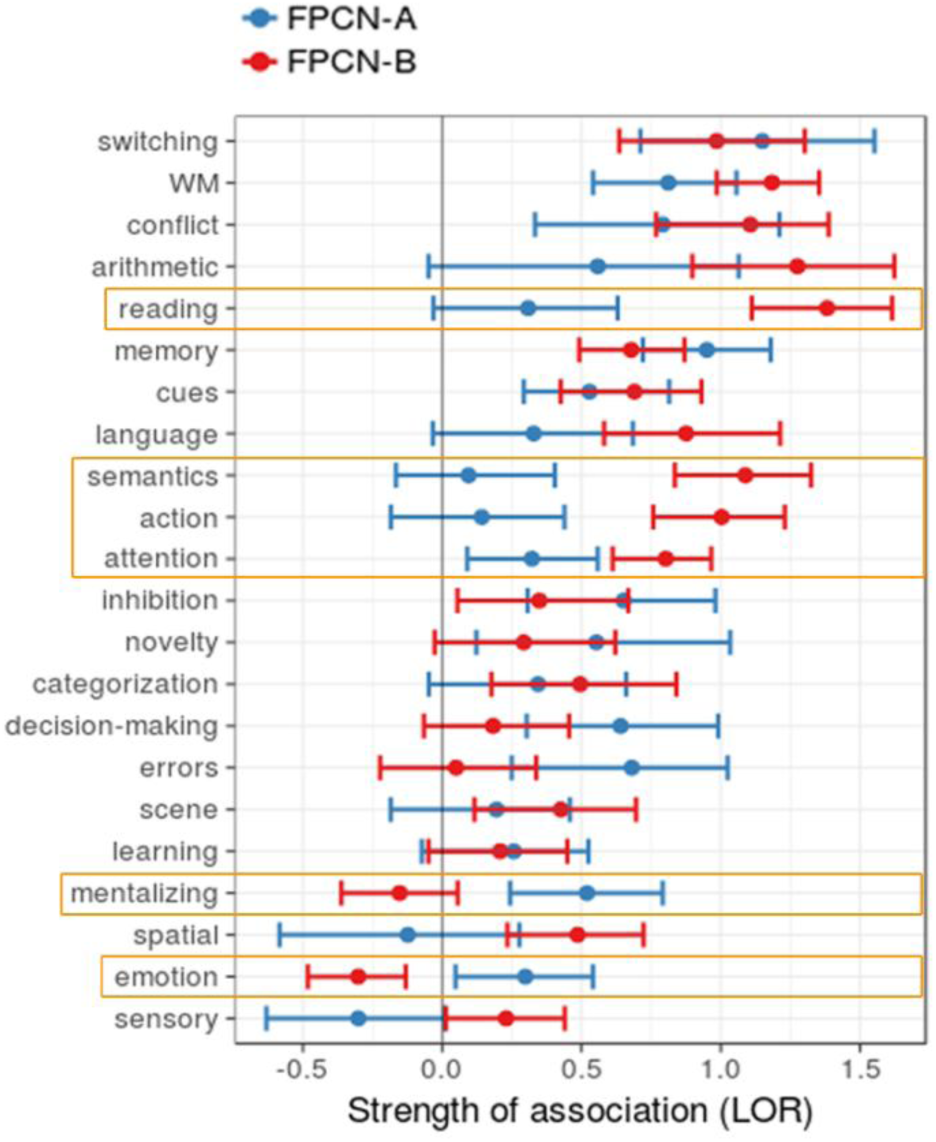
Meta-analytic functional preference profile of FPCN subsystems. We trained naïve Bayes classifiers to predict the presence or absence of activation in each FPCN subsystem using a set of 60 psychological topics and plotted topics that were significantly positively associated with at least one subsystem. Strength of association is measured in log odds-ratio (LOR) with values greater than 0 indicating that the presence of that topic in a study positively predicts activity in a subsystem. Ninety-five percent confidence intervals derived using bootstrapping are indicated, and topics differentially associated with each system are highlighted in orange.

## DISCUSSION

The current study provides novel evidence of highly reliable heterogeneity within the FPCN that is related to connectional patterns and functions associated with the DN and DAN―large-scale systems that contribute to internally-oriented thought and visuospatial perceptual attention, respectively. To summarize: (i) hierarchical clustering revealed a clear separation of FPCN_A_ and FPCN_B_ nodes based on intra-modular connections and inter-modular connections with the DN and DAN; (ii) a linear SVM classifier was able to distinguish FPCN_A_ and FPCN_B_ FC patterns with remarkable accuracy; (iii) differential coupling patterns were replicated in three additional data sets; (iv) Neurosynth meta-analytic coactivation patterns revealed differential task-based coactivation with the DN and DAN; (v) dynamic network interactions revealed that the FPCN fractionation relates to temporal variation in DN and DAN clustering patterns; and (vi) there are differences in the task domains that predict activation in FPCN_A_ and FPCN_B_. These findings place the organization and functions of the FPCN within the broader context of the brain’s network architecture, and offer a novel perspective on the systems-level network circuitry underlying executive control.

### Functional organization of the FPCN

Brain networks can be understood within the context of a hierarchical gradient of processing. At one extreme unimodal sensorimotor regions process concrete sensory and action-related information, while at the other extreme heteromodal regions elaborate upon such information, allowing for abstract thought, reasoning, and mental simulations of events^14-16^. Recently, it has been shown that the DN occupies a position that is further removed from sensorimotor processing than the DAN^14^. Our findings suggest that this distinction may be carried forward into the organization of the FPCN. We found that both FPCN subsystems were associated with topics related to core executive functions (e.g., working memory, conflict). However, FPCN_A_ demonstrated a relative bias towards the DN in terms of connectivity and functions, whereas FPCN_B_ demonstrated a relative bias towards the DAN in terms of connectivity and functions.

The DAN is activated when attention is directed in a top-down manner to task-relevant objects and locations, and also when intrinsically salient stimuli are detected^21-23,25,27,57,58^. Our findings suggest a close relationship between FPCN_B_ and the DAN in the network topology. Moreover, we found that FPCN_B_ was associated with functional domains that are known to activate the DAN. Specifically, FPCN_B_ was significantly more associated with topics related to attention and action than FPCN_A_. Prior work suggests that FPCN_B_ contributes to cognitive control by flexibly encoding task-relevant information including task rules (e.g., stimulus-response mappings) and their relationship to expected reward outcomes^2-4,6,59,60^. Notably, FPCN_B_ regions including the inferior frontal junction play a causal top-down role in modulating the DAN and perceptual attention^61,62^. One possibility is that FPCN_B_ represents information about task context in working memory and that the DAN translates this information into commands to guide the deployment of spatial attention to specific objects and locations^61,62^. By exerting top-down control over the DAN, FPCN_B_ may ensure that attention remains focused on task-relevant perceptual information, rather than salient, yet irrelevant stimuli, or task-irrelevant thoughts. Thus, the role of FPCN_B_ in executive control may be related to the abstraction, monitoring, and manipulation of sensorimotor contingencies to facilitate moment-to-moment interactions with the environment.

In contrast, FPCN_A_ regions are activated when attention is directed towards one’s own thoughts and away from perceptual inputs^40,63,64^, for example, during tasks that require meta-cognitive awareness^63,65,66^, relational reasoning^67^, multi-tasking and complex task sets^43,60,68-70^, stimulus-independent and abstract thinking^34,41,64,71-73^, mentalizing^74^, episodic memory^55,75^, future planning^5^, and prospective memory^76^. Consistent with this, we found that FPCN_A_ was preferentially coupled with the DN, which plays a role in bringing conceptual/associative knowledge to bear on current thought and perception^28-30,37,77^. Additionally, FPCN_A_ was associated with functional domains that are known to activate the DN. Specifically, FPCN_A_ was significantly more associated with topics related to mentalizing and emotion than FPCN_B_. Thus, FPCN_A_ may preferentially contribute to executive control in the context of introspective processes, and enable modes of thought that are relatively free from the constraints of concrete sensorimotor interactions with the environment. A recent framework^37^ suggests that FPCN_A_ (in particular the rostrolateral prefrontal cortex), may contribute to the abstract “top-level management” of thought, exerting a general constraint that keeps one’s focus on task-relevant material, yet allowing for some degree of spontaneous variability in thought. In this way, FPCN_A_ may play a role in regulating internal thoughts and emotions in service of social reasoning, mental time travel (e.g., future goal planning), and metacognitive awareness of emotional states. It may also contribute to the performance of traditional cognitive control tasks by allowing representations of abstract task rules and temporally extended contexts to guide the implementation of more concrete rules and actions^42,43,78,79^.

In every condition, including demanding cognitive control tasks (rule use, Stroop, 2-Back), we found robust coupling between FPCN_A_ and the DN. Consistent with this, a recent study found encoding of task-relevant information by the DN and increased activation during demanding rule switches, suggesting that it may contribute to some forms of cognitive control that involve activating different cognitive contexts^80^. We did find, however, that the magnitude of FPCN_A_-DN coupling was reduced during the cognitive control tasks relative to other conditions, and was significantly lower than FPCN_A_-FPCN_B_ coupling. FPCN_A_ was strongly aligned with the DN across the six conditions in the primary data set which were designed to elicit mental states that resemble those frequently experienced in everyday life. Thus, the diminished relationship with the DN during the traditional cognitive control tasks may represent the exception rather than the rule. FPCN_A_ may typically operate as an extension of the DN, but become co-opted by FPCN_B_ when it is necessary to perform highly complex perceptually-focused tasks. Thus, while FPCN_B_ may have evolved as an extension of the DAN processing stream to allow for the regulation of visuospatial perception and action during physical interactions with the environment (e.g., tool use), FPCN_A_ may have evolved as an extension of the DN processing stream to allow for the regulation of introspective processes such as complex social reasoning. This proposal aligns with suggestion that there is an intimate relationship between brain evolution, including expansion of the anterior prefrontal cortex in humans^81^, and the emergence of complex social life^82^. However, the functional distinction suggested here is just a starting point; a more elaborate theoretical framework will be required as work on the FPCN progresses.

### Relation to other models of executive control and frontoparietal organization

According to one model, the FPCN is critical for trial-by-trial adjustments in control, whereas a cingulo-opercular network is critical for the maintenance of task-goals across trials, supporting a balance between flexibility and stability^1^. Rapid adjustments in control may occur via flexible task-dependent shifts in FPCN coupling patterns^5,7,55^. Another model suggests that the “salience” network initiates shifts in modes of information processing related to the FPCN and DN^83^. Our findings suggest an orthogonal dimension of executive control, with different zones within the FPCN involved in visuospatial attention and more abstract thought processes, respectively. This division of labor between subsystems of the FPCN is broadly compatible with the idea of a rostro-caudal gradient in the lateral PFC based on abstractness of representational content^40-46^. Here, we extend this idea to the large-scale network organization within which the lateral PFC is embedded. Recent work suggests a “distance from sensory-motor processing” organizational principle, with more complex and abstract processing occurring in regions that are physically remote from primary sensory and motor cortices^14,16^. Our findings suggest that FPCN_A_ may be further removed from sensory-motor processing than FPCN_B_. Consistent with this, we observed that FPCN_A_ but not FPCN_B_ nodes were negatively correlated with primary sensory-motor regions (**Supplementary Fig. 5**). Thus, a general principle of functional organization may apply across different brain networks^14^ and within the FPCN itself.

Other work has emphasized that the FPCN is a flexible hub that coordinates processing across other networks in a task-dependent manner^5,7,84^. In the current study, we found that although FPCN_A_ and FPCN_B_ were aligned with the DN and DAN, respectively, there was evidence that FC patterns flexibly adapted to task demands. There were overall shifts in FPCN_A_ and FPCN_B_ coupling patterns, as well as shifts in the relative “preference” of coupling with the DN versus DAN. Thus, while FPCN_A_ can be distinguished from FPCN_B_, this reflects a *relative* and flexible difference in FC patterns rather than an absolute and fixed aspect of network architecture. The organization noted here is thus fully compatible with findings of task-dependent reconfiguration of FPCN FC patterns. Interestingly, we observed a weaker “selectivity index” for FPCN_B_, yet greater task-related flexibility across the task conditions examined here. This suggests that FPCN_B_ may have access to information from both the DN and DAN, and may be positioned to flexibly mediate interactions between more concrete visuospatial and more abstract information. It could be the case that task-related flexibility was lower for FPCN_A_ because it is exclusively interconnected with multimodal regions that process highly abstract information, and may not be equipped to interact with many regions. However, a different battery of tasks could potentially reveal greater FC flexibility in FPCN_A_ than FPCN_B_.

### Limitations

One of the challenges in examining heterogeneity within the FPCN it how to define this network to begin with. Rather than select a single method, we used nodes based on three different parcellations (Yeo, Gordon, and Power) that identified the FPCN as a functional unit on the level of other functional systems (e.g., visual and somatomotor networks). We then looked for finer-grained heterogeneity within this system. Notably, although the FPCN was defined slightly differently across parcellations, we observed a fractionation in each case. A second issue is that our range of tasks was not exhaustive making it possible that different network interactions could be observed in some contexts (e.g., positive coupling between the FPCN_A_ and DAN). One instance may be perceptual metacognition, which is known to rely on parts of the FPCN_A_ including the rostrolateral prefrontal cortex^65^. Additionally, it is possible that the FPCN may not fractionate, but rather, function as a domain general resource during demanding tasks that require considerable effort^4^. However, our findings do suggest that a FPCN fractionation can be observed in many diverse contexts. Finally, our analysis is limited by the reliance on pre-defined network boundaries and the assumption of discrete brain clusters/networks. Any brain parcellation is a dimensionality reduction on a complex space, and should be viewed as a general guiding principle rather than a set of fixed and precise brain network demarcations. Moreover, the network affiliation of a given brain region can shift across time and context^56,85^. That being said, our results provide evidence that spatially distinct parts of the FPCN―as defined using three different parcellations―are differentially coupled with the DN and DAN across a range of contexts.

### Conclusions

Executive control processes are multifaceted and likely rely on multiple interacting, yet distinct neural systems. The current work makes a step forward in discerning the network basis of executive control and may offer new predictions about clinical deficits in control functions. For example, altered connectivity between the FPCN_A_ and DN may interfere with regulating abstract information (e.g., self-referential thoughts) in conditions such as depression, whereas altered connectivity between the FPCN_B_ and DAN may interfere with regulating visuospatial attention (e.g., focusing on goal-relevant objects) in conditions such as ADHD. More broadly, our results are consistent with the notion that brain network organziation may, in part, reflect a gradient of representational abstraction.

## METHODS

### Participants

Participants in the primary data set were 24 healthy adults (Mean age = 30.33, SD = 4.80; 10 female; 22 right handed), with no history of head trauma or psychological conditions. The sample size was chosen based on norms in fMRI research. The ability to detect true effects was assessed using independent data sets (see below for details). This study was approved by the UBC clinical research ethics board, and all participants provided written informed consent, and received payment ($20/hour) for their participation. Due to a technical error, data for the movie and acceptance-based introspection conditions were not collected for one participant. At the end of scanning, one participant reported experiencing physical discomfort throughout the scan. Similar results were obtained with or without inclusion of this participant’s data, so they were included in the final analysis.

### Experimental task conditions

The primary data set included six ecologically valid task conditions in separate six-minute fMRI runs. Each task condition was designed to elicit a continuous mental state and did not require any responses. (1) *Resting state.* Participants lay in the scanner with their eyes closed and were instructed to relax and stay awake, and to allow their thoughts to flow naturally. (2) *Movie watching*. Participants watched a clip from the movie *Star Wars: Return of the Jedi* and were instructed to pay attention to the actions of the characters, and also to what they may be thinking and feeling. (3) *Artwork analysis*. Participants viewed four pieces of artwork in the scanner, each for 90 seconds. These pieces were pre-selected by participants, and during scanning, they were instructed to pay attention to the perceptual details of the art, their inner experience (i.e., thoughts and feelings), and what each image meant to them personally. (4) *Shopping task*. While in the scanner, participants viewed a pre-recorded video shot from a first-person perspective of items within several stores in a shopping mall. They were told to imagine themselves going through the mall in order to find a birthday gift for a friend, and to analyze each in terms of whether it would be a suitable birthday gift based on the preferences of their friend. *(5) Evaluation-based introspection*. Participants were asked to think about a mildly upsetting issue involving a specific person in their life (e.g., a friend, roommate, sibling, or partner), and asked to reflect on what the person and situation means to them, what has happened in the past and may happen in the future, and to analyze everything that is good or bad about the situation. *(6) Acceptance-based introspection*. Participants were asked to reflect on the same upsetting issue as in the previous case, but this time were instructed to focus on moment-to-moment viscero-somatic sensation, and to accept these sensations without any judgment or elaborative mental analysis.

### fMRI data acquisition

fMRI data were collected using a 3.0-Tesla Philips Intera MRI scanner (Best, Netherlands) with an 8-channel phased array head coil with parallel imaging capability (SENSE). Head movement was restricted using foam padding around the head. T2*-weighted functional images were acquired parallel to the anterior commissure/posterior commissure (AC/PC) line using a single shot gradient echo-planar sequence (repetition time, TR = 2 s; TE = 30 ms; flip angle, FA = 90°; field of view, FOV = 240 mm; matrix size = 80 × 80; SENSE factor = 1.0). Thirty-six interleaved axial slices covering the whole brain were acquired (3-mm thick with 1-mm skip). Each session was six minutes in length, during which 180 functional volumes were acquired. Data collected during the first 4 TRs were discarded to allow for T1 equilibration effects. Before functional imaging, a high resolution T1-weighted structural image was acquired (170 axial slices; TR = 7.7 ms; TE = 3.6 ms; FOV = 256 mm; matrix size = 256 × 256; voxel size = 1 x 1 x 1 mm; FA = 8°). Total scan time was ∼ 60 minutes. Head motion was minimized using a pillow, and scanner noise was minimized with earplugs.

### Preprocessing

Image preprocessing and analysis were conducted with Statistical Parametric Mapping (SPM8, University College London, London, UK; http://www.fil.ion.ucl.ac.uk/spm/software/spm8). The time-series data were slice-time corrected (to the middle slice), realigned to the first volume to correct for between-scan motion (using a 6 parameter rigid body transformation), and coregistered with the T1-weighted structural image. The T1 image was bias-corrected and segmented using template (ICBM) tissue probability maps for gray/white matter and CSF. Parameters obtained from this step were subsequently applied to the functional (re-sampled to 3 mm^3^ voxels) and structural (re-sampled to 1 mm^3^ voxels) data during normalization to MNI space. The data were spatially-smoothed using an 8-mm^3^ full-width at half-maximum Gaussian kernel to reduce the impact of inter-subject variability in brain anatomy.

To address the spurious correlations in resting-state networks caused by head motion, we identified problematic time points during the scan using Artifact Detection Tools (ART, www.nitrc.org/projects/artifact_detect/). Images were specified as outliers according to the following criteria: translational head displacement greater than 0.5 mm from the previous frame, or rotational displacement greater than .02 radians from the previous frame, or global signal intensity > 4 standard deviations above the mean signal for that session. The mean number of identified outliers was 4.93 (range: 0 - 15) and did not differ across conditions (*F* < 1). Each participant had at least 5.3 minutes of non-outlier time points. Outlier images were not deleted from the time series, but rather, modeled in the first level general linear model (GLM) in order to keep intact the temporal structure of the data. Each outlier was represented by a single regressor in the GLM, with a 1 for the outlier time point and 0 elsewhere.

Using CONN software^49^, physiological and other spurious sources of noise were estimated and regressed out using the anatomical CompCor method ^86^. Global signal regression was not used due to fact that it mathematically introduces negative correlations (Murphy et al., 2009). The normalized anatomical image for each participant was segmented into white matter (WM), gray matter, and CSF masks using SPM8. To minimize partial voluming with gray matter, the WM and CSF masks were eroded by one voxel. The eroded WM and CSF masks were then used as noise ROIs. Signals from the WM and CSF noise ROIs were extracted from the unsmoothed functional volumes to avoid additional risk of contaminating WM and CSF signals with gray matter signals. The following nuisance variables were regressed out: three principal components of the signals from the WM and CSF noise ROIs; head motion parameters (three rotation and three translation parameters) along with their first-order temporal derivatives; each artifact outlier image; linear trends. A band-pass filter (0.009 Hz < *f* < 0.10 Hz) was simultaneously applied to the BOLD time series during this step.

### Definition of networks and nodes

The nodes for most of our analyses were anatomical regions of interest (ROIs) created by Yeo and colleagues^87,88^ based on their 17-network parcellation ^12^. The 17-network parcellation was split into a set of 114 cortical regions composed of roughly symmetric territories in the left and right hemispheres, and were defined in relation to network boundaries, sulcal patterns, and confidence maps. For each network, spatially connected regions were combined to form a single ROI, whereas spatially disconnected regions became separate ROIs. Vertices near between-network boundaries were peeled back. The current analysis focused on 37 ROIs spanning the DN specifically the DN Core subsystem^29^, DAN, and FPCN. For each participant, we extracted the mean timeseries from participants’ unsmoothed data, to minimize the chance of signal contamination across ROIs. The residual timeseries (following nuisance regression) for each ROI was used to compute condition-specific correlation matrices. Where noted, we also analyzed data using DN, DAN, and FPCN nodes from the Gordon and Power parcellations, using the same preprocessing methods. For the analysis with the Gordon nodes (obtained from: http://www.nil.wustl.edu/labs/petersen/Resources.html), we derived the network structure empirically by submitting the group averaged correlation matrix (333 x 333 nodes) during rest to the Louvain community detection algorithm as implemented by the Brain Connectivity Toolbox^48^. The algorithm was run 1000 times, with the resolution parameter, gamma, set to 2.1. We then we computed a co-classification matrix reflecting the probability that each pair of nodes were assigned to the same module across iterations. We then ran the community detection algorithm on this co-classification matrix to determine the final modular structure, which resulted in a 9-network parcellation with 139 nodes spanning our networks of interest. For the analysis with the Power nodes, we used the nodes corresponding to DN, DAN, and FPCN in their network parcellation^53^ (94 ROIs; 5mm spheres created around MNI space coordinates, obtained from: http://www.jonathanpower.net/2011-neuron-bigbrain.html).

### Hierarchical clustering analysis

We used hierarchical clustering to organize nodes into a tree structure based on the similarity of their FC profiles, such that the nodes within each cluster are as similar as possible, and different clusters are as dissimilar as possible. We first created a group-averaged correlation matrix reflecting mean FC across all six conditions in the primary data set. We then extracted the subgraph composed of within-FPCN FC values and the subgraph composed of FPCN connections with the DN and DAN. These subgraphs were submitted to the hierarchical clustering algorithm (Cluster v3.0, 1988, Stanford University) which used the average linkage method to cluster nodes. In this method, the distance between two clusters is defined as the average distance between each point in one cluster to every point in the other cluster. Spearman correlation was used to determine distance. Cluster graphs were viewed with Java TreeView (v1.1.6r4 http://jtreeview.sourceforge.net).

### SVM classification analysis

We used a support vector machine (SVM) classifier to discern whether differences in FPCN_A_ and FPCN_B_ FC patterns were generalizable across participants. This analysis used nodes from the Yeo parcellation, and FPCN nodes were labeled as subsystem A or B based on the results of the intra-modular hierarchical clustering analysis. This analysis was not designed to test the strength of group level results with respect to FPCN FC patterns; rather it was designed to test consistency across participants (i.e., the extent to which FC patterns in a subset of participants would generalize to a different subset). The SVM classifier was implemented with RapidMiner software^89^. The cost parameter, C, was set to 1, and the convergence epsilon was set to .001. Our main analysis used a linear kernel, however, we also report results with an ANOVA kernel to demonstrate the robustness of our results (**Supplementary Figs. 2 and 4**). For each individual, we created a vector consisting of FPCN_A_ correlations with all FPCN nodes (excluding self connections), and a vector consisting of FPCN_B_ correlations with all FPCN nodes (excluding self connections), for the intra-modular analysis. For the inter-modular analysis we created a vector consisting of correlations between FPCN_A_ nodes and DN and DAN nodes, and a vector consisting of correlations between FPCN_B_ nodes and DN and DAN nodes. We excluded interhemispheric correlations which are likely to reflect indirect functional interactions. Additionally, we did not include correlation values for one FPCN_A_ region (pre-SMA), so that the FPCN_A_ and FPCN_B_ FC vectors would be equal in length. The correlation vectors served as input features (73 for the intra-modular analysis, and 76 for the inter-modular analysis) and were assigned a value of 1 or −1 to specify the FPCN subsystem to which they belonged. We tested the accuracy of the classifier using 4-fold cross-validation. The data were split into 4 equal-sized groups, with 75% of the data used for training the classifier, and the left-out 25% used for testing the classifier. This process was repeated 4 times until every participant was used in the testing set once. Participants’ data could not appear in both the testing and training set in the same iteration, and we did not perform any type of iterative optimization or feature selection. Thus, our analysis method should minimize the chance of overfitting^90^. Notably, when performing the classification analysis 50 times with network labels randomly re-shuffled, mean classification accuracy was at chance level in every condition for the intra-modular analysis (Rest: 50.5%; Movie: 50.0%; Artwork: 49.4%; Shopping: 49.1%; Evaluation: 49.3%; Acceptance: 48.4%) and for the inter-network analysis (Rest: 50.7%; Movie: 51.2%; Artwork: 48.3%; Shopping: 49.4%; Evaluation: 47.4%; Acceptance: 49.6%).

### Network visualization

For each task, the group averaged FC matrix was thresholded to retain connections with z(*r*) > .15, and then submitted to the Kamada–Kawai energy algorithm, implemented in Pajek software. This algorithm produces spring-embedded layouts that minimize the geometric distances of nodes based on their topological distances in the graph. Well-connected nodes are pulled towards each other, whereas weakly-connected nodes are pushed apart in a manner that minimizes the total energy of the system.

### Comparing mean between-network FC

After Fisher r-to-z transforming the correlation values, we averaged the z(r) values reflecting pairwise connections between the frontoparietal networks and the DAN and the DN, using Yeo parcellation nodes. We calculated average FC separately for the left and right hemispheres, and then collapsed across hemisphere, given the lack of statistical difference (i.e., there was no effect of hemisphere within any condition; all *P*'s > .05).

### Replication analyses

In all cases, data were analyzed using the same preprocessing methods as noted earlier. *Rule-based cognitive control task*. This data set (N = 15) has been described in full elsewhere^3^. Participants used one of two rules (male/female face discrimination or abstract/concrete word meaning discrimination) to respond to visual stimuli on each trial. On some trials subjects could earn money by responding quickly and accurately. The rules switched from trial to trial requiring participants to actively represent and flexibly switch between the different rules. Data from a single run (run 1 of 6) were analyzed. *Stroop Task*. This data set (N = 28) was acquired from the OpenfMRI database (accession number ds000164)^91^. Participants performed the color-word version of the Stroop task with three conditions (congruent, incongruent, and neutral) and were instructed to ignore the meaning of the printed word and respond to the ink color in which the word was printed. Data were acquired in a single run. *N-Back working memory task*. This data set (N = 41) was acquired from the OpenfMRI database (accession number ds000115)^92^. We analyzed the data from the task period during the 2-Back block in control participants. The task was to determine whether each letter was the same as the letter shown two trials previously.

### Neurosynth meta-analytic analyses

For the following two analyses, we used Neurosynth (www.github.com/neurosynth/neurosynth-data; version 0.6)―a diverse meta-analytic database of 11,406 fMRI studies spanning a wide range of published neuroimaging studies. Each study in the database is recorded as a set of peak activations in MNI coordinates, in addition to normalized frequencies of each word used in the abstract. Jupyter notebooks with analysis code and data are available at: https://github.com/adelavega/fpcn_fractionation. The analysis methods used here are similar to those introduced in two previous publications^93,94^, albeit with masks of each FPCN subsystem based on the Yeo 17-network parcellation, which was replicated in our data set using hierarchical clustering on the FPCN nodes.

#### Meta-analytic co-activation maps

To identify differential co-activation patterns across the brain between the two FPCN sub-networks, we performed a meta-analytic contrast between studies that activated FPCN_A_ and studies that activated FPCN_B._ The resulting images identify regions of the brain that were more likely to co-activate with one subsystem or the other. To determine statistical significance, we performed a two-way χ^2^test, and used the False Discovery Rate (*q* < 0.05) to threshold the resulting images. For display purposes, the resulting images were binarized and visualized using the pysurfer Python library (https://pysurfer.github.io/).

#### Meta-analytic functional preference profiles

We generated multivariate functional profiles for each FPCN subsystem by determining which psychological functions best predicted activity for each. We employed a set of 60 topics—which concisely and robustly represent the semantic information in Neurosynth—derived using latent dirichlet allocation (LDA) topic-modeling^95,96^. For each FPCN subsystem, we selected a set of studies that activated at least 5% of voxels within its mask, and a set of studies that did not. We then trained a naives Bayes classifier to differentiate the two sets of studies based on loading of each topic onto each study in the database. From the fitted classifiers, we extracted the log odds-ratio (LOR) of a topic being present in activate studies versus inactive studies. In other words, we calculated how useful each semantic topic was in determining if a study reported activation within a given FPCN subsystem. We determined the 95% confidence interval of the LOR for each topic using bootstrapping, by sampling with replacement from the full set of studies 1000 times. This allowed us to determine (in a post-hoc, exploratory analysis) if topics were significantly associated with each network, and if these associations differed between FPCN_A_ and FPCN_B_.

### Task-related flexibility

The flexibility index was computed as the similarity of FC patterns *within* a given condition (from the first half to the second half) minus the similarity of FC patterns *between* conditions. A larger difference implies that FC patterns changed more across than within conditions, implying task-related flexibility. For each participant and condition, we extracted and vectorized FC values (Fisher transformed correlation values) reflecting FPCN_A_ correlations with DN and DAN nodes, and values reflecting FPCN_B_ correlations with DN and DAN nodes. We used the Pearson correlation as a measure of the similarity of the FC vectors for each pair of conditions. These correlation values were Fisher transformed and averaged, to arrive at a single value reflecting the similarity of FC across conditions for the FPCN_A_ and a single similarity value for the FPCN_B_. We additionally computed the similarity of FC patterns within each condition from the first half (first three minutes) to the second half (last three minutes) of each condition. By subtracting between-condition from within-condition similarity values, this provides a selective measure of the effect of condition on FC patterns―our measure of flexibility. Notably, this is a conservative estimate of flexibility given that less data was used in calculating within-context FC values, likely resulting in less reliable and lower similarity values. To determine the flexibility of FC for each FPCN ROI, we computed the variability (SD) of FC across contexts with each DN and DAN node, and then averaged across these values.

### Dynamic FC analysis

We conducted a novel dynamic FC analysis to examine the possibility of a FPCN fractionation within the context of dynamic network interactions. Prior work has shown that functionally-relevant connectivity patterns can be isolated from ∼ 60 seconds of data. To examine time-varying connectivity patterns, the data were filtered (0.0167 Hz < *f* < 0.10 Hz) based on the window size of 60-seconds in order to limit the possibility of detecting spurious temporal fluctuations in FC^97^. Within each window, we computed the mean strength of between-network FC for each pair of networks, thus providing time-series of between-network FC values, and we also computed the mean weighted clustering coefficient for the DN and DAN, providing a time-series of clustering values. The weighted clustering coefficient, *C*^*W*^, quantifies the potential for communication within the immediate neighborhood of a node, and is defined as the proportion of neighbors around node *i* that are also interconnected, normalized by the average “intensity” of connections. It is computed as:

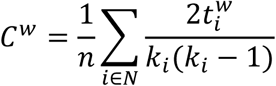

where 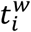 is the number of triangles around node *i* (i.e., a set of three nodes that are all interconnected), normalized by their intensity (edge weight), and *k*_*i*_ is the degree (total number of connections) of node *i*. We modified the computation of the weighted clustering coefficient so that only *within-network* nodes were considered. This served two purposes: (i) it allowed for a meaningful interpretation of resulting clustering values; and (ii) the standard clustering coefficient would have resulted in an artificial correlation between dynamic interactions and changes in clustering strength. For example, because DN nodes are connected with FPCN_A_ nodes, they would normally be included in the computation of the clustering coefficient for each DN node. However, this would create a spurious positive correlation between DN clustering and FPCN_A_-DN FC strength. Thus, to compute our modified weighted clustering coefficient, we extracted relevant within-network connections; set self-connections and negative FC values to 0; normalized all connections by the strongest weight (from the set of all matrices) such that weight magnitudes were rescaled to the range [0,1]; computed the weighted clustering coefficient for each node using the Brain Connectivity Toolbox; averaged clustering values across nodes within a given network. Mean weighted clustering was computed for each window. For each participant, we then calculated the correlation between the time-series of between-network FC values and clustering coefficient values. We then fisher r-to-z transformed these correlations to allow for statistical testing at the group level.

### Statistical Analyses

For all analyses, reported P-values were two-tailed and corrected for multiple comparisons as noted. In the machine learning analyses we emphasized classification accuracy which is easily interpreted since baseline accuracy was 50% (with a balanced class size), and did not compute statistical significance. To test for differential coupling patterns, we submitted mean FC values to repeated measures ANOVAs and follow-up paired t-tests with P-values that corrected for the number of comparisons (6 with respect to the DN and 6 with respect to the DAN). Accordingly, P-values reported as statistically significant at P < .05 Bonferroni corrected were below the threshold of P = .008. For the coactivation analysis, images were whole-brain corrected using a false discovery rate of *q* = 0.05. For the dynamic FC analysis, we used paired t-tests to compare FPCN_A_ and FPCN_B_ relationships. In this case there were 9 comparisons relevant to the DN and 9 comparisons relevant to the DAN, so P-values reported as statistically significant at P < .05 Bonferroni corrected were below the threshold of P = .005. To examine flexibility across contexts, flexibility index values for each subsystem were assessed using one-sample t-tests relative to 0. To examine whether FPCN_A_ and FPCN_B_ were more coupled with each other or the DN/DAN we submitted FC values to paired t-tests (FPCN_A_-FPCN_B_ FC versus FPCN_A_-DN FC or FPCN_A_-FPCN_B_ FC versus FPCN_B_-DAN FC) and report P-values that were Bonferroni corrected for the 9 comparisons.

## Author contributions

MLD and KC developed the design of the study. MLD performed data collection. MLD, ADV, and CM conducted the analyses. JRA-H, RNS, MWC, and KC provided guidance on analysis methodology. KC supervised this research. MLD wrote a manuscript draft. All authors commented on the manuscript.

